# Phosphorylation of HspB1 regulates its mechanosensitive molecular chaperone interaction with native filamin C

**DOI:** 10.1101/325712

**Authors:** Miranda P. Collier, T. Reid Alderson, Carin P. de Villiers, Daisy Nicholls, Heidi Y. Gastall, Timothy M. Allison, Matteo T. Degiacomi, Dieter O. Fuerst, Peter F.M. van de Ven, Kristina Djinovic-Carugo, Andrew J. Baldwin, Hugh Watkins, Katja Gehmlich, Justin L.P. Benesch

**Affiliations:** Department of Chemistry, Physical and Theoretical Chemistry Laboratory, University of Oxford, South Parks Road, Oxford, OX1 3QZ, U.K.; Division of Cardiovascular Medicine, Radcliffe Department of Medicine and British Heart Foundation Centre of Research Excellence Oxford, University of Oxford, Headington, Oxford, OX3 9DU, U.K.; Biomolecular Interaction Centre and School of Physical and Chemical Sciences, University of Canterbury, Christchurch, 8140, New Zealand; Department of Chemistry, Durham University, South Road, Durham, DH1 3LE, U.K.; Department of Molecular Cell Biology, Institute for Cell Biology, University of Bonn, D53121 Bonn, Germany; Department of Structural and Computational Biology, Max F. Perutz Laboratories, University of Vienna, Campus Vienna Biocenter 5, A-1030 Vienna, Austria; Department of Biochemistry, Faculty of Chemistry and Chemical Technology, University of Ljubljana, Večna pot 113, SI-1000 Ljubljana, Slovenia

**Keywords:** HspB1/Hsp27, chaperone, phosphorylation, mechanosensing, filamin, catch-bond, mass spectrometry, ion mobility, X-ray crystallography, NMR spectroscopy

## Abstract

Small heat-shock proteins (sHsps; HspBs) are molecular chaperones involved in the cellular stress response and a range of basal functions. Despite a multitude of targets, sHsp interactions are not well understood due their heterogeneous structures and weak binding affinities. The most widely expressed human sHsp, HspB1, is prevalent in striated muscle, where the actin cross-linker filamin C (FLNC, γ-filamin, ABP-L) is a putative binding partner. Musculoskeletal HspB1 is phosphorylated in response to a variety of cues, including mechanical stress, which promotes oligomer disassembly and association with myoarchitectural elements. Here, we report the up-regulation and interaction of both proteins in the hearts of a mouse model of heart failure, with HspB1 being phosphorylated and FLNC increasingly associated with the sarcomeric Z-disc. We used a combination of structural approaches to reveal that phosphorylation of HspB1 results in increased availability of the residues surrounding the phosphosite, facilitating their interaction with folded FLNC domains equivalent to a force-sensing region in the paralog filamin A. By employing native mass spectrometry, we show that domains 18 to 21 of FLNC are extensible under conditions mimicking force, with phosphorylated HspB1 stabilising an intermediate from further unfolding. These findings report on conformations accessible during the cycles of mechanical extension central to filamin function, and are consistent with an interaction between the chaperone and a native target that is strengthened upon the application of force. This may represent a new mode of molecular chaperone activity, allowing HspB1 to protect FLNC from over-extension during mechanical stress.

## Introduction

Mechanosensitive proteins withstand cycles of force-induced structural changes, and as such possess a pliable ‘native’ state. This is critical for their function, whether to provide elasticity to cells and tissues, open and close a channel in the membrane, or reveal cryptic or force-enhanced binding sites in mechanotransductive signalling pathways (1, 2). Excessive force, however, can lead to loss of function due to unfolding and subsequent formation of deleterious aggregates that overburden the capacity of the protein quality control network (3, 4). Such pathological deposits of intracellular proteins underlie a variety of muscle myopathies (5, 6). To counter the effects of misfolding in general, cells invest in molecular chaperones. The frequency of association between one class of chaperone in particular, the ATP-independent small heat-shock proteins (sHsps, HspBs] (7), with the cytoskeletal support network (8–11) and muscle contractile machinery (12–14) suggests a role for mechanosensitivity in sHsp-client binding.

One potential such interaction is between HspB1 (also known as Hsp27 in humans) and filamin C (FLNC; also known as γ-FLN; ABP-L; FLN2). Filamins, of which there are three paralogs in adult humans, are large modular proteins that play key roles in signalling pathways, cytoskeletal organization, cellular motility and differentiation (15, 16). They cross-link cytoskeletal actin into branched three-dimensional networks and are components of adhesion assemblies. FLNC, almost exclusively found in striated muscle, associates with thin filaments of sarcomeric actin at the myofibrillar Z- and intercalated discs (17–19) (Fig. 1A).

**Figure 1.**
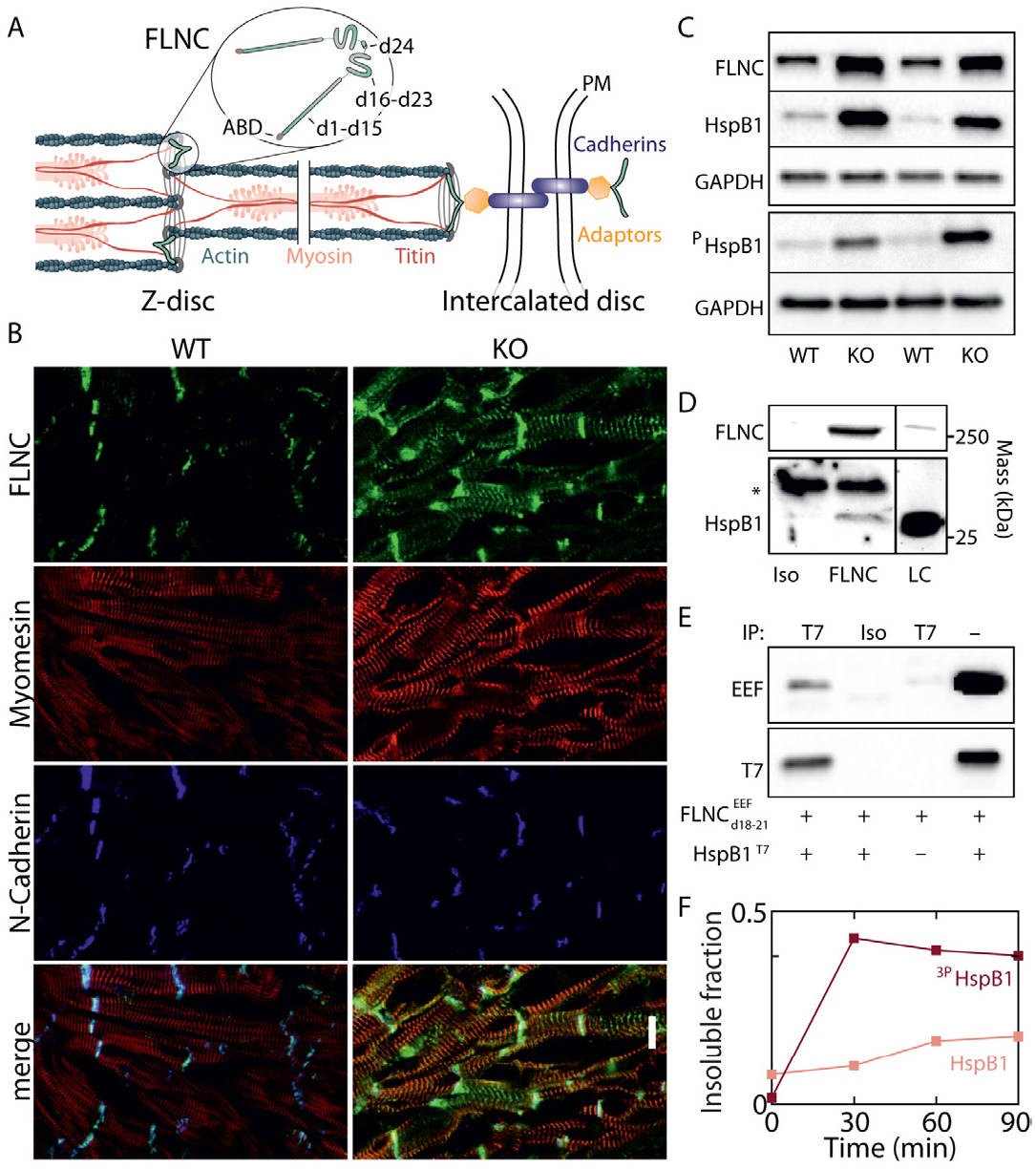
FLNC and HspB1 are up-regulated in MLP KO mouse heart and interact via domains 18-21 in a phosphorylation-dependent manner. **A** Schematic of FLNC composition and localization in striated muscle at the Z-disc and intercalated disc. Grey lines = multiple Z-disc proteins, including α-actinin, desmin, etc. Adaptors = talin, vinculin, catenins, etc. PM = plasma membrane. Not pictured: possible localization at costameres. **B** Frozen sections of ventricular tissue from WT and MLP KO mice. Sections were stained for FLNC (green) and counter-stained to visualize sarcomeres (myomesin, red) and intercalated discs (N-cadherin, blue). Scale bar = 10 μM. Merged images show FLNC is primarily localised at intercalated discs and sarcomeric Z-discs, and is more abundant in the DCM tissue. **C** Blots for FLNC, HspB1, and ^pS86^HspB1 in WT and MLP KO mouse hearts, and for GADPH as a loading control. Both FLNC and HspB1 were up-regulated, and HspB1 was phosphorylated at Ser86 (equivalent to human Ser82). **D** Immunoprecipitation from MLP KO ventricular tissue lysate using an FLNC-specific antibody or isotype control; and loading control. Molecular masses from positions of markers, * indicates signal from antibody chains. HspB1 co-precipitates with FLNC, evidencing an interaction between the endogenous proteins. **E** Recombinant EEF-tagged FLNCd18-21 mixed with T7-taggedHspB1 and immunoprecipitated. T7 antibody (but not an isotype control) pulled down both HspB1 and FLNC_d18-21_, indicating binding between the recombinant proteins. Neither protein is pulled down when HspB1-T7 is omitted from the sample. **F** Pelleted fraction of a 1 to 2 mixture of recombinant FLNC_d18-21_ and WT or ^3D^HspB1 over the course of 90 minute incubation at 25°C. Measurements by densitometry of SDS-PAGE (Fig. S1).

Alterations in the human *FLNC* gene can lead to severely impaired myoarchitecture and large protein aggregate deposits, and were initially linked to skeletal myopathies (designated ‘filaminopathies’), with some carriers also displaying cardiac abnormalities (20–24). Recently, the range of known disease-linked *FLNC* alleles has expanded to include many associated with cardiac pathologies, often without skeletal manifestation. These encode variants implicated in hypertrophic cardiomyopathy (HCM), restrictive cardiomyopathy (RCM), and dilated cardiomyopathy (DCM) (27–33). HspB1 was found to be prevalent in aggregates from patients with filaminopathy (25) and at sarcomeric lesions with FLNC (26), suggesting a role for their possible association during stress.

Human filamins are ≈280-kDa homodimers, each monomer consisting of an N-terminal actin-binding domain followed by 24 immunoglobulin-like (Ig) domains (d1-d24), the last of which mediates dimerization (Fig. 1A) (34). The distal region (dl6-d24) encompasses the binding sites for most filamin partners (16), which appears to include HspB1, since this interaction was first identified in yeast two-hybrid assays using the segment d19-d21 of FLNC. In filamin A (FLNA), the distal region can sense mechanical cues by partially extending under force to reveal cryptic binding sites (35–38). FLNC contains a large insertion within d20, and less is known about its structure and the molecular basis of its roles. Evidence that FLNC can sense local force includes: functionality that necessitates a degree of plasticity in response to tension (39); 73% sequence identity and overlapping roles and binding partners with FLNA (40, 41); the ability to protect myofibrillar structural integrity (42); and differential in-cell labelling of shielded cysteines with applied force (43).

Each HspB1 monomer is composed of a conserved core α-crystallin domain (ACD) flanked by a short, flexible C-terminal region, which makes intra-oligomer contacts (45), and a longer N-terminal domain (NTD). The NTD also confers oligomeric stabilization (46, 47), with phosphorylation at Ser15, Ser78 or Ser82 causing a shift from a large, polydisperse ensemble to smaller oligomeric species, including dimers (48–50). HspB1 is phosphorylated at Ser82 (the most prevalently modified site (51)) in response to mechanical cues, inducing translocation to the sarcomere (52) and sites of increased traction force within the cytoskeleton (53, 54), a mechano-accumulation phenomenon similarly observed for FLNC (2, 55). Candidate substrates in these areas also include actin and overstretched titin; however, upon HspB1 phosphorylation its affinities to these are reported as diminished (56) and unaltered (57), respectively, suggesting that phosphorylation may trigger or modulate the interaction between HspB1 and FLNC in particular.

To investigate the impact of phosphorylation on HspB1 intra-oligomer contacts and its interaction with FLNC, we employed experimental approaches spanning the tissue to atomic levels. We validated the proposed interaction between FLNC and HspB1 and noted up-regulation of both proteins, including the phosphorylated form of HspB1, in a mouse model of heart failure. We found that a segment of the NTD just downstream of the ACD in HspB1 adopts a range of conformations that are modulated by phosphorylation to trigger oligomer dissociation. A peptide segment from this flexible region binds multiple FLNC Ig domains that we show adopt an extensible conformation. Performing Coulombically driven unfolding in the gas phase as a proxy for the mechanical forces FLNC experiences *in vivo* allowed us to probe features of the interaction that are invisible at rest. This revealed that phosphorylated HspB1 inhibits a partially extended form of FLNC from further unfolding. Our results support a catch-binding mode between a molecular chaperone and a mechanosensitive target that is modulated by phosphorylation, and may be important for cardiac function upon myoarchitectural perturbation.

## Results

### FLNC and HspB1 are up-regulated and interact in DCM hearts

We first sought to test the association between HspB1 and FLNC in cardiac tissue. To examine the effect of impaired cardiac biomechanics on FLNC expression and localization, we used muscle LIM protein (MLP) knock-out (KO) mice. MLP, like FLNC, is specific to striated muscle, and the phenotype of these mice reproduces the morphology of DCM and heart failure in humans (58). Frozen sections of ventricular tissue from wild-type (WT) and MLP KO mice were stained for FLNC, and counter-stained to visualize sarcomeric structure (myomesin) and intercalated discs (N-cadherin) (Fig. 1B). FLNC was located at intercalated discs and Z-discs, at greater abundance in MLP KO hearts based on fluorescence intensity, and with no apparent aggregation. This is consistent with previously reported FLNC localization (l9) and with the role of intercalated discs and Z-discs as force-bearing sites (59). Western blotting, using lysates from WT and MLP KO mouse hearts, confirmed that FLNC was up-regulated in the diseased tissue relative to WT (Fig. 1C). HspB1 was up-regulated in the same samples, including its phosphorylated form (at S86, analogous to S82 in humans; mouse HspB1 does not contain a phosphosite equivalent to human S78 (60)) (Fig. 1C). A proportion of HspB1 was co-immunoprecipitated together with FLNC from lysates of MLP KO ventricular tissue, indicating an interaction between these two endogenously expressed proteins (Fig. 1D).

Next we expressed EEF-tagged human FLNC domains 18-21, encompassing the apparent HspB1 binding site (39), and full-length human HspB1 including a T7 tag. Pulling down the HspB1 protein with the T7 antibody co-immunoprecipitated FLNC_d18-21_ (Fig. 1E), confirming a direct interaction between these proteins. To investigate the effect of HspB1 phosphorylation on the interaction, we expressed WT human HspB1 and ^3P^HspB1, a triple mutant containing phosphomimetic Ser to Asp substitutions at residues 15, 78, and 82. Upon incubation with FLNC_d18-21_ *in vitro*, we noticed that mixtures accumulated insoluble protein aggregates while isolated proteins remained soluble. In a 2:1 mixture of the chaperone to the FLNC fragment, both proteins increasingly migrated to the pelleted fraction over time, an effect that was substantially increased by phosphomimicry (Fig. 1F, Fig. S1). The results support an interaction between HspB1 and FLNC_d18-21_ to form large complexes, that can be modulated by phosphorylation of HspB1.

### Phosphomimicry reduces oligomer size and coincides with disorder in the N-terminal a region of HspB1

To interrogate the regulatory role of phosphorylation, we modified a previous construct of the structured ACD (HspB1_84-171_) (45) to incorporate a small part of the N-terminal domain just upstream of the β2 strand, which we term the “principal phosphorylation region” (PPR) as it includes phosphosites Ser78 and Ser82 (HspB1_77-m_). We also mutated these sites to mimic phosphorylation events (^2P^HspB1_77-m_). Two-dimensional nuclear magnetic resonance (NMR) spectra of each of the three constructs featured well-dispersed resonances (Fig. 2A, Fig. S2A), evidencing folded dimer structures. The HspB1_84-171_ and HspB1_77-171_ spectra overlay closely, with additional peaks in the region diagnostic of disorder (7.8-8.6 ppm in the ^1^H dimension) arising in the longer construct from the additional N-terminal residues (Fig. 2A). To examine the impact of phosphorylation on the PPR, we compared the NMR spectra of HspB1_77-171_ and ^2P^HspB1_77-171_. We noted clear differences, particularly intensified signal in the disordered region for the phosphomimic, consistent with an increase in flexibility (Fig. 2A, Fig. S2A). Chemical-shift-based order parameters and secondary structure propensities indicated a high degree of backbone flexibility and a low amount of secondary structure for the β2 strand and the PPR (Fig. 2B).

**Figure 2.**
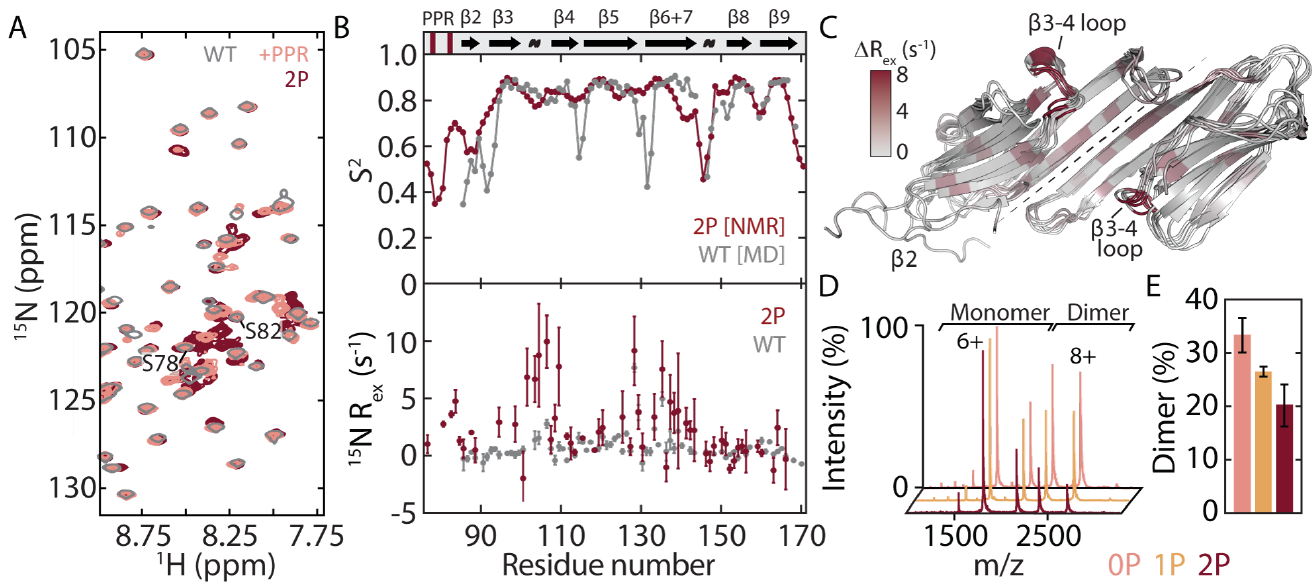
HspB1_77-171_ undergoes dynamical changes leading to partial dissociation upon phosphomimicry. **A** Overlaid ^1^H-^15^N HSQC spectra of HspB1_84-171_ (gray), HspB1_77-171_ (pink), and ^2P^HspB1_77-171_ (maroon). More signals are present in the region 7.8-8.6 ppm upon phosphomimicry. **B** N-H order parameters (*S*^*2*^) derived using assigned backbone chemical shifts from ^2P^HspB1_77-171_ (maroon) and from MD simulations of HspB1_84-171_ (gray) confirm that the PPR and β2 strand are highly dynamic (upper). A plot of *R*_ex_, a qualitative metric of μs-ms motions probed by ^15^N CPMG relaxation dispersion, for HspB1_84-171_ (gray) and ^2P^HspB1_77-171_ (maroon) reveals a significant increase in dynamics in residues 95-110 (lower). Secondary structure elements from PDB 4MJH are indicated. **C** Snapshots from a 1-μs MD simulation of HspB1_84-172_ showing the complete dissociation of a β2 strand. Residues are colored according to *R*_ex_ (from **B**), showing the region of increased dynamics to be in the β3-β4 loop on the side of the ACD. **D,E** Native mass spectra and quantification of HspB1_77-171_ WT (pink), IP (^P^Ser82, yellow), and 2P (^P^Ser78-^P^Ser82, maroon). The construct populates a monomer-dimer equilibrium, and phosphomimicry shifts the oligomeric state toward the monomer.

To visualise the structural fluctuations of β2, we performed a 1-μs molecular dynamics (MD) simulation of the HspB1 ACD starting from a structure (PDB: 4MJH) in which the β2-strand is intra-molecularly hydrogen bonded to β3. We found that over the course of the simulation β2 fluctuated considerably, readily losing its secondary structure and detaching from the rest of the ACD (Fig. 2C). Calculation of the order parameters from the simulation reveals good correspondence with that from our NMR data (Fig 2B), and is consistent with the disorder observed by others (61).

We next looked at chemical shift perturbations (CSPs) in the HspB1 ACD upon phosphomimicry. These were widely dispersed but small in magnitude (Fig. S2B), supporting similarity of the ACD structures. To probe for effects on internal dynamics, we recorded ^15^N spin relaxation measurements that monitor both ps-ns and qs-ms motions. Whereas the fast dynamics were unaffected (Fig. S2C), residues 95-110 (including the loop between β3 and β4) in the phosphomimic displayed evidence of conformational exchange on the slower timescale (Fig. 2B; Fig. S2D-E). We reasoned that this could indicate transient sampling of a lowly populated state involving contact between the PPR and the ACD. The major state would then contain an unbound PPR, consistent with the small CSPs between HspB1_84-171_ and HspB1_77-171_ for residues 102-110, lack of any conformational exchange slow enough to give rise to multiple peaks, and flexibility of the N-terminal residues in solution (Fig. 2B).

Absence of significant broadening in the NMR spectra as a function of increasing HspB1_77-171_ concentration suggested that the contacts made by the PPR regulate intra- rather than inter-dimer binding. To test this hypothesis, we recorded mass spectra under conditions preserving noncovalent interactions (62). The data revealed that HspB1_77-171_ populates a monomer-dimer equilibrium that is shifted toward dissociation upon incorporation of one and two phosphomimics (Fig. 2D-E). The absence of higher-order oligomers indicates that appreciable inter-molecular binding of the PPR is unlikely in the context of this truncated construct. The combined NMR and MS data are therefore consistent with phosphorylation modulating an order-to-disorder transition by β2 and the PPR that leads to oligomer dissociation via rearrangement of intra-dimer contacts.

### The PPR of HspB1 interacts with a pocket in the ACD

We next carried out crystallographic trials, employing a strategy of mixing peptide mimics of the PPR, varying in length and degree of phosphorylation, with a well-ordered ACD construct (HspB1_84-170_). This led to a structure of HspB1_84-170_ bound to ALSRQL^P^SSGVSEI (residues 76-88 phosphorylated at Ser82), solved by molecular replacement in space group P12_1_1, that contains four dimers in the asymmetric unit (Table S1, Fig. S3A). The monomers within each dimer differ at their N-terminal ends: the β2 strand of one is intra-molecularly hydrogen bonded to its own β3 strand, whereas in the other these residues are extended and stabilized by contacts with a β4-β8 groove of a neighboring dimer (Fig. 3A-B). This groove also acts as a binding pocket for an intra-subunit interface within the C-terminal region (Fig. S3B) (45).

**Figure 3.**
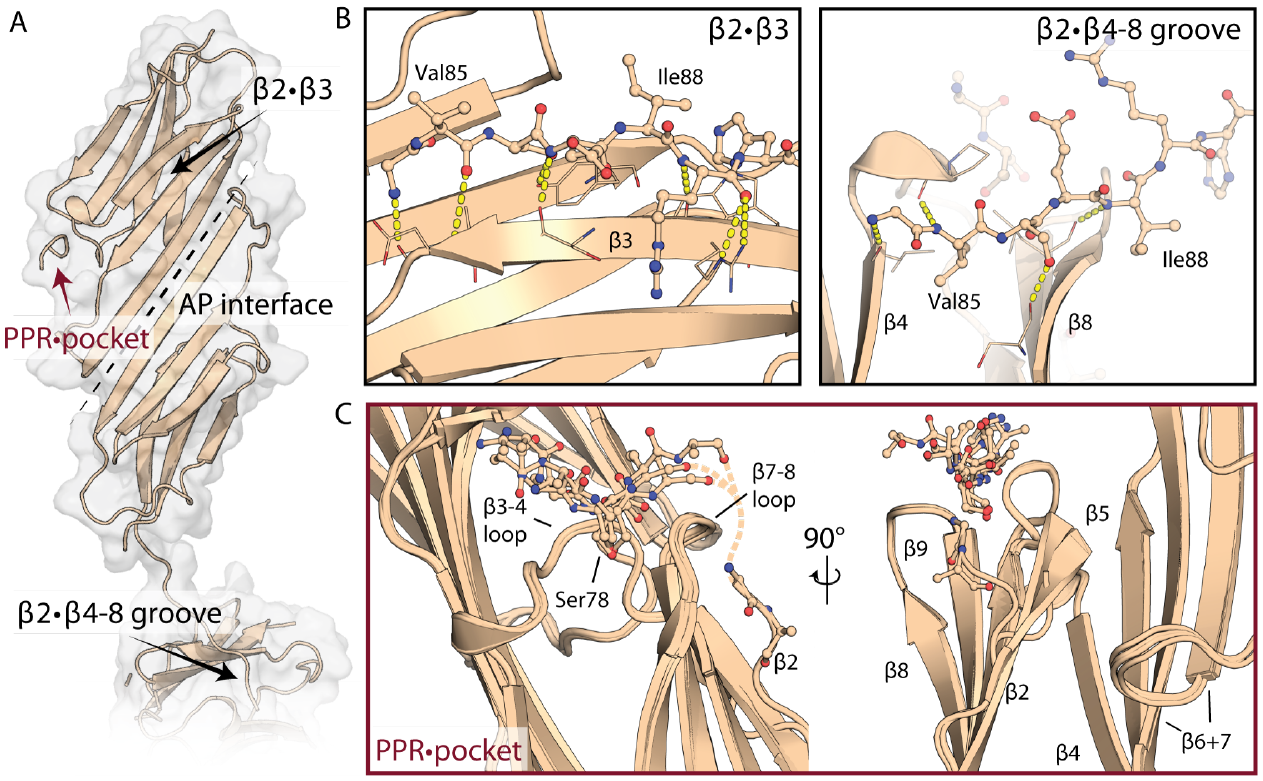
Crystal structure of the HspB1 ACD in complex with a partial peptide mimic of the PPR. **A** HspB1 ACD dimer with peptide PPR mimic bound. Dotted line indicates the antiparallel inter-dimer interface. **B** Areas indicated by black arrows in *A* showing two locations of the β2 strand: bound to β3 (left) or dissociated from the ACD and reaching across the β4-β8 groove of a neighboring monomer (right). **C** PPR binding (red arrow in **A**), shown as an overlay of the four peptide-bound monomers in the ASU. Whereas the ACDs overlay closely, the peptides appear heterogeneous and flexible. Rotated view shows β2 strand divergences, sometimes arcing toward the peptide binding site and in one instance partially modelled as continuation of the peptide.

Upon refinement of the ACD, the 2Fo-Fc and difference density maps displayed a clear unmodeled feature occupying a pocket between the β3-β4 and β8-β9 loops, which lies on the side of the subunit close to the dimer interface. This feature was only present adjacent to subunits with intra-molecularly bound β2 strands (Fig. S3C), and aligns with the dynamical changes we observed by NMR in the β3-β4 loop (Fig. 2B-C). We were able to build peptides into this density, but only accommodating subsets of the 11 possible residues, with their side-chain densities remaining poor in contrast to the ACD (see Methods). An ensemble of the four peptide-bound ACDs provides the best overview of the interaction available from these data (Fig. 3C). Our results reveal how the β2 strand and PPR assume heterogeneous conformations. Though predominately disordered in solution (Fig. 2B-C), this region is also capable of making intramonomer associations with the β3 strand and stretching to the pocket between β3-β4 and β8-β9 loops (Fig. 3B-C), as well as inter-subunit interactions by occupying a β4-β8 groove in a neighboring dimer (Fig. 3B). While it is not clear what the relative populations of these states are in the context of the full-length HspB1, our combined data reveal that this heterogeneity is modulated by phosphorylation, resulting in changes in the stabilities of the interfaces that are likely responsible for shifting the equilibrium away from large oligomers towards smaller species.

### HspB1 residues 80-88 bind multiple domains within FLNC

To isolate a stable HspB1-FLNC_d18-21_ complex and localize the binding within HspB1, we performed native ion mobility mass spectrometry (IM-MS) experiments (63). FLNC_d18-21_ displayed an envelope of peaks centered on low charge states, and a collision cross-section (CCS) of 37.2 ± 0.8 nm^2^ (Fig. S4A). This is consistent with a compact rather than a linear architecture (Fig. S4B), suggesting an inherently extensible conformation for this FLNC segment, wherein noncovalent inter-domain interactions would be susceptible to rupture under force. Next, based on the HspB1 sites whose exposure shifts upon phosphorylation, we mixed FLNC_d18-21_ with either HspB1_84-171_ or peptides mimicking the PPR, since this site was phosphorylated in the heart failure model mice (Fig. 1C). We observed charge state distributions corresponding to complexes between FLNC_d18-21_ and one or two copies of the peptide HspB1_80-88_ (or its phosphorylated counterpart ^P^HspB1_80-88_) (Fig. S5). Complexes were not observed in mixtures of FLNC_d18-21_ with HspB1_84-171_ or with negative controls (Fig. S5), revealing a specific interaction with this N-terminal segment of HspB1.

To measure the association of ^P^HspB1_80-88_ and HspB1_80-88_ to FLNC_d18-21_, we recorded mass spectra at increasing peptide concentrations (Fig. 4A, Fig. S6). By quantifying the relative abundances of FLNC_d18-21_ and the peptide-bound species, we could determine binding affinities (Fig. 4A, lower panel; Fig. S6C). We obtained similar dissociation constants (*K*_D_S) of 130 ± 17 μM (HspB1_80-88_) and 110 ± 10 μM (^P^HspB1_80-88_), indicating weak binding that is not affected substantially by phosphorylation. These findings contrast with the notable effect of HspB1 phosphomimicry on the interaction of full-length chaperone with the FLNC fragment (Fig. 1E). However, they can be rationalised by phosphorylation increasing the effective concentration of the binding region by modulating its conformational state (Fig. 2B, Fig. S2) and leading to dimer (Fig. 2D) and oligomer disassembly (48–50). To localise binding of HspB1_80-88_ to domains with Ig folds or to the d20 insertion (which is possibly disordered), we prepared a modified FLNC construct lacking the 82 insertion residues (FLNCd_18-21,ΔI_). This construct also binds multiple HspB1 peptides (Fig. S5C), indicating that HspB1 recognizes folded Ig domains of FLNC.

**Figure 4.**
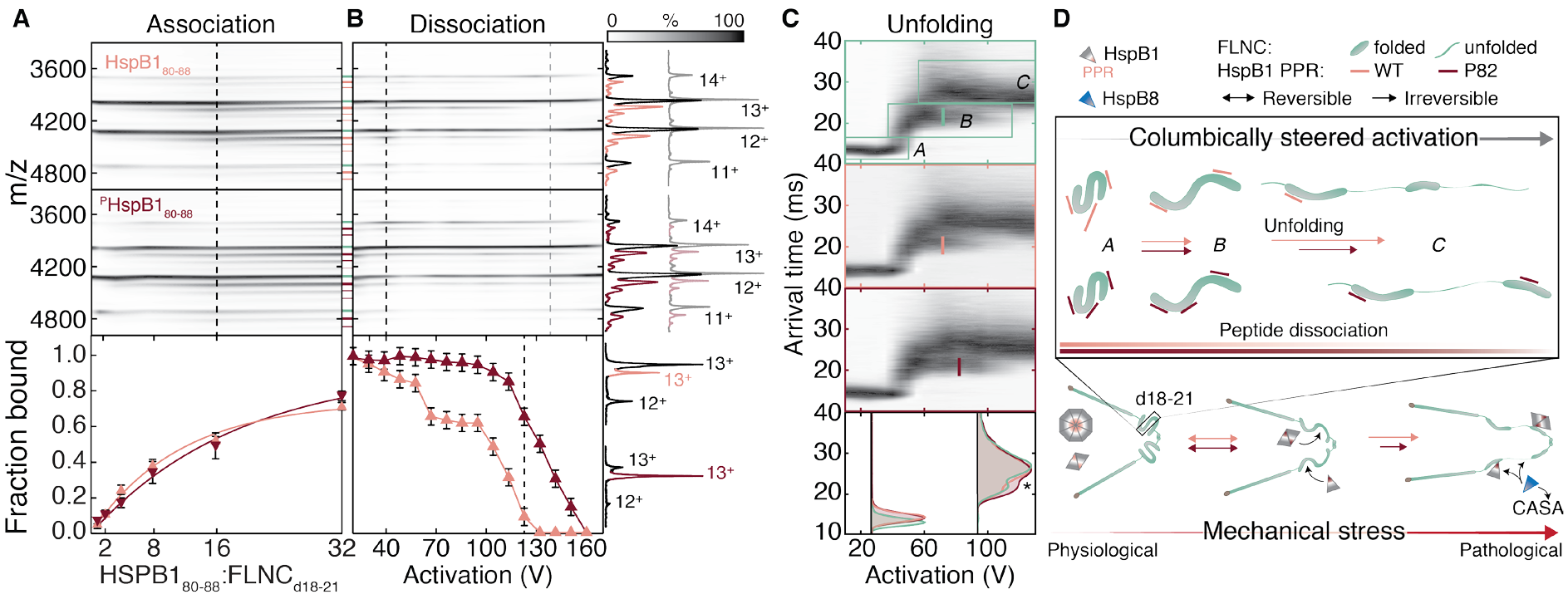
HspB1_80-88_ binds FLNC_d18-21_ with phosphorylation modifying the force-accessible behaviour of the complex. **A** Contour plots of native mass spectra of FLNC_d18-21_ obtained upon titrating the unphosphorylated (upper) and phosphorylated (middle) peptides. Association of multiple peptides can clearly be observed (pink lines between plots; same line thickness designates the same charge series and hence the same stoichiometry). Quantification of the relative abundances allows the determination of binding curves, revealing similar affinities of each peptide to FLNC_d18-21_ in solution (lower). **B** Dissociation of the peptide was affected by activation in the gas phase, with the phosphorylated peptide remaining bound to higher potentials than the unphosphorylated equivalent. Breakdown curves (derived from tandem MS data, projections to the right and Fig. S7) allow quantification of this effect, and reveal clear differences between the peptides. Mass spectra matching dashed lines in **B** are projected on the right. At 40 V, binding differences are negligible, but at 140 V, all of the unphosphorylated peptide has been ejected but the phosphopeptide has not. Error bars represent one standard deviation of the mean. **C** Coulombically steered unfolding of the 13+ charge stage of FLNC_d18-21_ unbound, bound to one HspB1_80-88_, and bound to one ^P^HspB1_80-88_. Bottom, overlaid arrival time distributions at 30 V and 90 V, highlighting the persistence of the intermediate state with bound phosphopeptide (*). Lines on unfolding plots designate the activation required to transition half of the intermediate to the final state. **D** Schematic of force-induced changes to FLNC underlying the unfolding plots, and relation to HspB1 peptide binding. Both peptides bind multiple sites (Ig domains) within resting, compact FLNC_d18-21_. During extension, inter-domain interactions are disrupted without peptide ejection, and eventually individual domains unfold, attenuated by the phosphopeptide. The lower panel places the findings in the context of mechanical stress and HspB1 oligomerization. At low stress, the majority of unmodified HspB1 self-associates and the PPR is minimally exposed. At the onset of mechanical stress, HspB1 is phosphorylated rapidly, whereupon it dissociates to reveal the binding site and can recognize extending FLNC. ^P^HspB1 then inhibits later stages of unfolding, which may be irreversible. Under extreme stress, CASA is triggered, involving recruitment of HspB8.

### Phosphorylation modifies the force-accessible trajectory of the FLNC-HspB1 complex

Having found marginal difference in binding affinities between the phosphorylated and unphosphorylated HspB1 peptides with FLNC_d18-21_ at equilibrium, we next sought to probe the strength of this interaction under the application of force. The motivation for this stemmed from the observation that the filamin distal region (Fig. 1A) undergoes extension *in vivo* (43), with FLNA domains even predicted to be able to unfold fully prior to breakage of the dimer interface, at physiologically relevant forces and loading rates (1, 39). As a proxy for mechanical stress, we performed gas-phase experiments in which protein ions are activated by collisions with inert gas, leading to conformational perturbations and Coulombic-repulsion-driven chain extension (64–66). Providing an analogous endpoint to mechanical unfolding, an advantage of this approach is that, by capitalising on the sensitivity and capability of ion selection in the mass spectrometer, even low-affinity complexes can be interrogated individually (67).

Upon extending the HspB1_80-88_-FLNC_d18-21_ complex by ramping the activation potential, we observed a gradual reduction in the abundance of peptide-bound FLNC_d18-21_, indicating dissociation of the peptide (Fig. 4B). Strikingly, phosphorylation of HspB1_80-88_ increased greatly the energy required to remove the peptide from FLNC: at 140 V, all unphosphorylated HspB1_80-88_ had dissociated, while ^P^HspB1_80-88_-bound FLNC persisted until 180 V (Fig. 4B, right). To quantify this differential dissociation behaviour, while avoiding potential cross-talk between charge states (68), we selected the 13+ ion of FLNC_d18-21_ bound to a single peptide in the mass spectrometer, and performed analogous activation ramps (Fig. S7). This allowed us extract the fraction of FLNC_d18-21_ associated with each peptide as a function of activation (Fig. 4B, lower). The breakdown curve for the FLNC_d18-21_:HspB1_80-88_ complex revealed two inflection points, at ≈55 V and ≈110 V. This suggests that HspB1_80-88_ associates with FLNC_d18-21_ in (at least) two locations, consistent with the observation of multiple peptides binding in our titration experiments (Fig. 4A), ^p^HspB1_80-88_ dissociated less readily from FLNC_d18-21_, and the breakdown curve features only one inflection point at ≈130 V. This suggests that phosphorylation increases both the selectivity and strength of interaction of HspB1 with FLNC during extension.

In an attempt to rationalise the observation of phosphorylation-dependent differences under activation (Fig. 4B) but not in its absence (Fig. 4A), we examined the unfolding trajectory of FLNC_d18-21_. Ion mobility (IM) measurements revealed that, over the range of activation potentials that dissociate HspBi peptides from FLNC_d18-21_, it extends from the initial folded state (*A*) through an intermediate (*B*), to reach a final state (*C*) (Fig. 4C, upper). The same three states were observable for FLNC_d18-21_ with either peptide bound (Fig. 4C). In all cases, unfolding commenced at ≈40 V, a lower potential than necessary for peptide dissociation. This suggests that HspB1 does not bind exclusively the regions within FLNC_d18-21_ that extend first, since unfolding of a ligand-bound domain will coincide with ejection of the ligand (69). The stability of *A*, based on the potential required to transition to *B*, was unaffected by either peptide. *B*, however, required more activation to transition to *C* when bound to ^P^HspB1_80-88_ (≈75 V) versus HspB1_80-88_ (≈60 V) (Fig. 4C). This difference was consistent across charge states and instrument conditions (Fig. S8). The data points to the formation of interactions when the phosphopeptide-bound FLNC structure is activated, leading to its increased resistance to extension, and dissociation at higher potentials than the unmodified peptide. This suggests a chaperone-target mechanism reliant on the recognition of native-like states but stabilisation of unfolding intermediates, in a manner that is regulated by post-translational modification.

## Discussion

We have investigated how phosphorylation of HspB1 modulates its interaction with FLNC. Our biophysical and structural studies reveal conformational heterogeneity in the β2 strand and the immediately downstream PPR, which encompasses the primary HspB1 phosphosite. These residues are largely disordered in our construct in solution, but also make intra-molecular contacts with β3 and a pocket on the edge of the ACD in a manner that is regulated by phosphorylation. We observed an inter-dimer interaction between the β2 strand and the β4-β8 groove, also the location of a C-terminal inter-dimer interaction (45), in our crystal structure. This is consistent with alternative modes of occupancy for the β4-β8 groove observed in other vertebrate sHsps (70, 71), and hints at intra-molecular competition for the binding site in the context of the HspB1 oligomers. The conformational plasticity may be the primary source of the modulation in chaperone activity associated with HspB1 phosphorylation (50, 72–74), exposing binding sites both directly through intramolecular rearrangements, and indirectly by causing oligomeric dissociation (48–50).

Similar heterogeneity of N-terminal residues has been observed in other sHsps and shown to regulate assembly (75), including in HspB5 and HspB6, both of which co-assemble with HspB1 *in vivo* (76). The HspB5 β2 strand and residues immediately preceding it exist in multiple conformations (77), including bound to β3 both intra- (45, 71) and inter-monomer (77). Recent crystal structures of HspB6 show no resolved density for β2, and support disorder-to-order transitions upon interactions made by the N-terminus (78). In both HspB5 (79) and HspB6 (80), these contacts are controlled by phosphorylation. Taken together, these observations indicate a mechanism of tuneable structure and function via phosphorylation-based modulation of local flexibility. Changing the conformational landscape and binding properties of heterogeneous structures in this manner offers a way to balance entropic loss with enthalpic gain (81), potentially key for the ATP-independent chaperone activity of sHsps (82).

We observed that HspB1 associates with FLNC in the hearts of MLP KO mice, an established animal model of heart failure, with concomitant up-regulation of both proteins and phosphorylation of the chaperone. We replicated this interaction *in vitro*, and localised it to HspB1_80-88_, encompassing part of the PPR and β2 strand, which binds FLNC_d18-21_ at multiple locations. While phosphorylation of HspB1 increased the association by promoting unbinding and flexibility of the PPR, it does not require unfolding of the target. This behaviour may be facilitated by HspB1 and FLNC_d18-21_ both containing Ig folds, such that chaperone-target interfaces mimic intra-molecular interactions in HspB1. Such interaction between a sHsp and a target in its native state is one of few observed directly (78, 83), and contrasts with the canonical chaperone mode of recognizing unfolded or misfolded substrates (84).

We mimicked mechanical strain on FLNC by performing experiments in which we unfolded the protein in the gas phase, along a pathway defined by the Coulombic repulsion between mobile charges (66). While the free energies of unfolding transitions can be influenced by the lack of solvent in these experiments (85, 86), electric field-induced protein mechanics have been shown to reflect functional trajectories (87). Our experiments revealed that FLNC_d18-21_ adopts a compact state, meaning that multiple inter-domain interactions are likely. This coheres with predictions of its similarity to FLNA_d18-21_, whose semi-compact L-shape forms part of a propeller-like FLNA_d16-21_ region with β-strand-swapping between domain pairs (35, 88, 89). Upon extending FLNC_d18-21_, we observed a series of conformational changes. Notably, transitions occurred without ejection of bound ^P^HspB1_80-88_, and involved an unfolding intermediate stabilized by ^P^HspB1_80-88_ relative to HspB1_80-88_. The results are consistent with a phosphorylation-triggered “catch-bond” or “molecular clutch”: an interaction with a longer lifetime under the application of force than in its absence (59), suggesting that a function of HspB1 is to regulate the extension of FLNC.

The extent of FLNC extension *in vivo* depends on various interactions, including at its dimer and actin interfaces, since breakage of these would lessen the strain borne by the putative mechanosensing region (1). While quantification of these interactions has not been reported for FLNC, reversible FLNA domain unfolding and actin unbinding are energetically similar, and each more common than dimer rupture at physiological forces and low loading rates (90, 91). Higher loading rates increase rupture forces and therefore the likelihood of domain deformation (90). Since catch-bond lifetimes are loading-rate-dependent (92), the ^P^HspB1-FLNC interaction may be energetically favoured under conditions of stress. This concurs with the increase in expression of both proteins in hearts of MLP KO mice and, since these animals display severe defects in myofibrillar organization (58), implies functional importance in the maintenance of myoarchitectural integrity. FLNC may require chaperoning to function properly under stress and avoid adverse structural disruption. In this role, HspB1 binding would be preventative rather than a response to damage, lowering the risk of FLNC hyperextension.

Nevertheless, critical force seems to overburden this system, exposing aggregation-prone regions of filamin and triggering chaperone-assisted selective autophagy (CASA) to clear and degrade filamin in muscle tissue (93–95). CASA involves recognition of compromised filamin by HspB8, and subsequent scaffolding of HspB8 to Hsp70 via the co-chaperone BAG3. Additional sHsps may participate in autophagy, since all those that are expressed in muscle can bind BAG3 except HspB7 (93, 96), which interestingly binds FLNC (97). HspB1, however, is the only sHsp now known to bind native FLNC in addition to BAG3, and readily co-assemble with several sHsp paralogs. Its co-localization with FLNC and BAG3 at sarcomeric lesions (26), and ability to interact with both compact and extending filamin, mean HspB1 may be a key hub in the proteostasis of muscle both during physiological and to pathological conditions (Fig 4D).

Our results, in the context of these previous reports, point to a two-tier, sHsp-mediated proteostasis mechanism for maintaining FLNC and the myoarchitecture, active at varying tensions. This is modulated by the cellular response to mechanical stress, via HspB1 phosphorylation, exposure of its FLNC-binding-site, and stabilization of the reputed mechanosensitive region of FLNC. The findings add to recent reports of molecular chaperones binding to force-bearing proteins (99–101), and may provide a new avenue for understanding filaminopathies and FLNC-linked cardiac-specific diseases.

## Materials and Methods

### Co-immunoprecipitation

1 μg of T7-tagged purified recombinant HspBi protein was added to 10 μg EEF-tagged recombinant FLNC_d18-21_ in IP buffer (0.05% Triton X-100, 1% BSA, protease inhibitors [Roche mini complete ETDA-free tablets) in PBS) and incubated for 1h, shaking, at room temperature. This was followed by another 30 minute incubation, shaking at room temperature after the addition of 0.5 μg anti-T7 (Novagen, 69522) or mouse IgG2b isotype (Abcam, ab18457) antibody. Next, 25 μl Protein G sepharose resin (Generon, PC-G5) was added to each tube and incubated for 1 hour, shaking at room temperature. The beads were then washed 4 times with IP buffer, re-suspended in 20 μl 5X SDS sample buffer (312.5 mM Tris HCl pH 6.8, 500 mM DTT, 10% SDS, 30% Glycerol, 0.05% bromophenol blue), and boiled at 95°C for 5 minutes. Immunodetection was performed using the anti-T7 (Novagen, 69522) or anti-EEF/ anti-tubulin (YL1/2) (Abcam, ab6160) antibodies.

### Analysis of MLP KO mouse hearts

Experimental procedures were performed in accordance with the UK Home office guidelines (project licences 30/2444 and 30/2977) and approved by respective institutional review boards. Animals were housed in specific pathogen free conditions, with the only reported positives on health screening over the entire time course of these studies being for *Tritrichomonas sp* and *Entamoeba spp*. All animals were housed in social groups, provided with food and water ad-libitum, and maintained on a 12h light:12h dark cycle (150-200lux cool white LED light, measured at the cage floor). MLP knockout mice (58) were backcrossed with C57BL/6J for > 6 generations before generating homozygous MLP knockout mice. Wildtype C57BL/6J were obtained from Harlan. Animals (females, age 3 months) were sacrificed by cervical dislocation, hearts flushed with ice-cold PBS and snap frozen in liquid nitrogen.

Immuno-fluorescence from frozen tissue was performed as described (102) using FLNC antibody RR90 (103), pan-cadherin antibody (Sigma) and myomesin antibody B4 (DSHB, Developmental Studies Hybridoma Bank). Immunoblotting from total protein extracts was performed as described (102) using anti-HspB1 (Cell Signaling, 2442), anti-^P^HspB1 (Ser82) (Cell Signaling, 2401), anti-GAPDH (Merck, ABS16) and FLNC (R1899; MRC PPU Reagents and Services, University of Dundee) antibodies. Immunoprecipitation was performed as described (102) using 5 ug of FLNC (R1899) antibody. Blotting for HspB1 (Cell Signaling, 2442) was performed using “Veriblot” (ab131366) as secondary HRP conjugate.

### Protein expression and purification

A pET28d(+) vector encoding human HspB1 ACD (residues 84-171) was transformed into *E. coli* BL21(DE3) cells (Agilent), and expressed and purified as described previously (45). For crystallography, residue K171 was deleted using a site-directed mutagenesis kit (Agilent) and HspB1_84-170_ was expressed and purified as described. HspB1_77-171_ and variants thereof were cloned into the same vector and expressed in the same manner, with both proteins containing a residual Gly-Ser overhang between the TEV protease recognition site and the beginning of the HspB1 sequence. Phosphomimic mutations S78E and S82D were introduced using site-directed mutagenesis, expressed as described, and purified as follows: cells were lysed using a cell press with added protease inhibitor cocktail (Roche). Lysate was centrifuged at 20,000 g for 20 minutes and the supernatant filtered and loaded onto a HisTrap column in buffer 20 mM Tris, 150 mm NaCl, 20 mM imidazole, 5 mM BME, pH 8. Protein was eluted with a gradient to 500 mM imidazole, pooled, and dialyzed overnight with TEV protease into the loading buffer at room temperature. The sample was then exchanged into the same buffer with addition of 8 M urea by cycles of centrifugation and dilution using an Amicon concentrator with 3.5 kDa cut-off. The cleaved and unfolded protein was passed over a HisTrap column and flow-through collected and refolded by overnight dialysis at room temperature into 100 mM NaCl, 20 mM Tris, pH 7.5. Protein was then concentrated and DTT added to a final concentration of 1 mM for storage at −20°C until use. For NMR experiments, HspB1 variants were expressed in M9 minimal medium containing ^13^C-labeled D-glucose as the only carbon source and ^15^N-labeled ammonium chloride as the only nitrogen source, and purified as described, with a final SEC (Superdex 75, GE Healthcare) step for exchange into the NMR buffer. Concentrations were determined by using a BCA assay (Thermo Pierce).

Full-length human HspB1 (WT or 3D, containing mutations S15D, S78D, and S82D) encoded in pETHSUL vector with an N-terminal SUMO and 6xHis-tag was transformed into *E. coli* BL21(DE3) cells. Cells were used to inoculate a 2 mL LB culture and grown for 7 hours at 37°C. Next 5oo qL of this culture was transferred to 1 L of ZYP auto-induction media. Cells were left to grow overnight at 24°C and 180 rpm until reaching OD 4, and then another 24 hours at 18°C and 180 rpm. Cells were harvested and lysed using a cell press with added protease inhibitor cocktail, spun down at 20,000xg for 20 minutes, and loaded onto a HisTrap column in 50 mM sodium phosphate, 250 mM NaCl, 12.5 mM imidazole, 5 mM BME, pH 7.5. Protein was eluted with a gradient to 500 mM imidazole. Eluent was pooled, dialyzed overnight with SUMO hydrolase at 4°C into the loading buffer, and the sample then passed over a HisTrap column and flow-through collected. This was concentrated and further purified by size exclusion chromatography using a Superdex 200 16/600 column (GE Healthcare). Purity was assessed by SDS-PAGE and protein was concentrated, flash-frozen, and stored in PBS with 1 mM DTT at - 80°C. Concentration was determined by absorption at 280 nm.

The region of human FLNC comprising domains 18-21 (WT, residues 1943-2408; Ig-only, residues 1943-2408 excluding 2163-2243) was encoded in a modified pET23a/EEF (PMID:8830773) with C-terminal 6xHis and EEF tags. The plasmid was transformed into *E. coli* BL21(DE3) pLysS cells (Agilent). Cells were grown in LB at 37°C until an OD600 of 0.6-0.8 was reached. Isopropylthiogalactoside (IPTG) was then added to a final concentration of 0.5 mM to induce expression for 16-18 hours at 21°C. Cells were resuspended in 300 mM NaCl, 50 mm Tris, 20 mM imidazole, 5 mM BME, pH 7.7 containing protease inhibitor cocktail and lysed using a cell press. Lysate was clarified by centrifugation at 20,000 g for 20 minutes, filtered through a 0. 22 μm filter, and applied to a 5 mL HisTrap column (GE Healthcare). Protein was eluted with a gradient to 500 mM imidazole, pooled, concentrated using an Amicon centrifugal concentrator with 30 kDa MWCO, and further purified by size exclusion chromatography using a Superdex 200 (10/300) column (GE Healthcare) at 4°C. Purity was assessed by SDS-PAGE and protein was stored in 200 mM ammonium acetate pH 6.9 at −80°C and used for experiments within two weeks of purification, after noticing in initial preparations some propensity for destabilization and aggregation following long term storage.

Peptides were purchased from Biomatik at 99+% purity with acetylated N-termini when the N-terminal residue was Glu80. Lyophilized peptides were resuspended in ddH_2_O to a stock concentration of 15 mM, flash-frozen, and stored at −20°C until use.

### HspB1-phosphopeptide crystallization and structure determination

Crystals were obtained following initial screens using 18 mg mL^−1^ HspB1_84-170_ with a 4-fold molar excess of one of several peptide mimics of the N-terminal region varying in length and degree of phosphorylation. Several combinations yielded reproducible crystals, but most exhibited poor diffraction (>7 Å). Following rounds of optimization using hanging-drop plates (VDX; Hampton Research), we collected a dataset from a crystal grown from 5 mM peptide ^P82^HspB1_76-88_ and 16 mg mL^−1^ HspB1_84-170_ that allowed for structure determination (Table S1). Drops were set using a protein-peptide mixture in PBS pH 7.0 and equilibrated against 22% PEG 3350; 0.02 M sodium potassium diphosphate; and 0.1 M Bis-Tris propane pH 7.5. The protein was mixed 1:1 with the crystallization solution (0.9 μl each) and drops were left to equilibrate at room temperature against a 1 mL reservoir. Crystals grew within 2 days and continued growing to maximum size within 6 days. Crystals were cryo-protected by serial transfer from the drop to the crystallization solution containing additional 10% PEG 400 and 5% followed by 10% glycerol, before being flash-frozen in liquid N_2_. Diffraction data were collected under cryogenic conditions on beamline I04-1 at Diamond Light Source, Harwell, UK.

Diffraction data were integrated using iMosflm (104) and scaled and merged using Aimless in the CCP4 suite with truncation at 2.1 Å. Phases were determined by molecular replacement using Phaser (105) with a previous HspB1 X-ray structure input as a search model (PDB 4MJH, modified by removal of flexible loops, bound C-terminal peptide, and β2 strands). The four dimers in the asymmetric unit were completed through cycles of manual building in Coot (106) and refinement in Phenix (107) imposing non-crystallographic symmetry (NCS) restraints. Peptide density was then apparent in the 2Fo-FC and Fo-Fc maps next to one monomer within each of the four dimers. Peptide fragments were placed through cycles of manual building in Coot and refinement in Phenix, now without NCS restraints and with treatment of TLS parameters (108). Most peptide side chains remained irresolvable throughout cycles of refinement in contrast to the well-resolved core. In addition, the peptide density never extended beyond a length accommodating 3 to 6 of 11 residues.

Thus, we attempted to deduce the orientation and identity of the residues captured in the density. In light of data suggesting an intramolecular interaction, we treated each peptide as though extending toward the β2 strand of the monomer to which it is bound. The question of which residues occupy the binding pocket was thus restricted to those that could reach from the β2 strand if the chain were continuous, eliminating the latter half of the peptide and leaving, roughly, ALSRQL. The lack of density for the remaining half of peptide is unsurprising, since most of these residues are repeated in the ACD and would compete for contacts with their counterparts which are covalently bound to the core and exist at a much higher local concentration, leaving this portion of the peptide no site to bind and thus too disordered to give rise to observable electron density. In each peptide region, the clearest side chain electron density pointed into a pocket between Phe104 and Gly161 that could not accommodate a bulky residue, but must be larger than alanine. Backbone density extends two residues N-terminally from this pocket, excluding Leu77 and implicating Ser78 or Leu81. Modeling each of these and refining caused negligible difference in R_work_ and R_free_; so based on close examination of the resulting Fo-Fc maps and potential chemical contacts within the pocket, as well as the NMR data supporting dynamic changes in this region of the ACD upon Ser78 mutation, we placed Ser78 into this location. At each dimer interface, Cys137 density evidenced both disulphide-bonded and reduced conformations, and so we modelled each of these with occupancy 0.5 in all cases, consistent with the labile nature of this bond (45). The final model had R_work_/R_free_ 21%/25%. It was deposited with atomic coordinates and structure factors in the RCSB PDB with the ID code 6GJH.

### NMR spectroscopy

All NMR spectroscopy experiments were recorded at 298 K on a 14.1 T Varian Inova spectrometer equipped with a 5 mm z-axis gradient triple resonance room temperature probe. Spectra were processed with NMRPipe (109) and analyzed with NMRFAM-Sparky (110). 2D sensitivity-enhanced ^1^H-^15^N HSQC spectra were acquired with ^1^H (^15^N) 512 (64) complex points, spectral widths of 8012 Hz (1800 Hz), maximum acquisition times of 64 ms (35.6 ms), an interscan delay of 1 s, and four scans per FID for a total acquisition time of 19 minutes. ^13^C,^15^N-^2P^HspB1_77-171_ was prepared at a final concentration of 0.9 mM in 30 mM sodium phosphate, 2 mM EDTA, pH 7 for resonance assignments, wherein HNCA, HNCO, and HN(CA)CO spectra were recorded. The assigned ^1^H^N^, ^15^N, ^13^CO, and ^13^Cα chemical shifts were analyzed with TALOS-N (111) and RCI (112) to respectively estimate the secondary structure and N-H order parameters.

Protonated, ^15^N labelled proteins were prepared at 0.9 mM (^2P^HspB1_77-171_) or 1 mM (HspB1_84-171_) for ^15^N relaxation studies. Standard pulse sequences to measure transverse relaxation times (*T*_2_), ^15^N heteronuclear nuclear Overhauser enhancements (hetNOE) (113), and ^15^N CPMG relaxation dispersion (114) were employed. The *T*_2_ experiments contained eight delay times up to 154 ms and an inter-scan delay of 2 s. The intensity changes over time (I/I_0_) were fit to an exponential decay, and values are reported as rates (i.e. 1/*T*_2_) for convenience. hetNOE experiments contained an inter-scan delay of 8 s, with values representing the intensity upon amide proton saturation divided by the intensity in its absence. CPMG relaxation dispersion experiments were recorded with variable delays between 180° pulses in the CPMG pulse train (τ_CPMG_ = 4*v*_CPMG_^−1^). A constant relaxation delay of 39 ms for the CPMG period and twenty *V*_CPMG_ values ranging from 54 to 950 Hz were employed. Peak shapes were fit with FuDA (115) to extract peak intensities, which were then converted into *R*_2,eff_ values using the following relation: *R*_2,eff_(*V*_CPMG_) = −1/*T*_*relax*_ ln(*I*(*v*_*CPMG*_)/*I*(0)), where *I*(*V*_*CPMG*_) is the intensity of a peak at *V*_*CPMG*_, *T*_*relax*_ is the constant relaxation delay of 39 ms that was absent in the reference spectrum, and *I*(0) is the intensity of a peak in the reference spectrum. Two duplicate *v*_*CPMG*_ points were recorded in each dispersion data set for error analysis, and uncertainties in *R*_2,eff_ were calculated using the standard deviation of peak intensities from such duplicate measurements. From plots of *R*_*2,eff*_ as a function of *V*_*CPMG*_, *R*_*ex*_ was determined by taking the difference of *R*_2,eff_ (54 Hz) and *R*_2,eff_ (950 Hz) with error bars representing the propagated errors in *R*_2,eff_.

### Native IM-MS

Because HspB1 can form disulphide-linked dimers, experiments involving the ACD were performed in the presence of 10-fold molar excess DTT. Gold-plated capillaries were prepared in-house. Native MS and IM-MS data were recorded on a Synapt G1 mass spectrometer (Waters) modified for the transmission of intact noncovalent protein complexes (116). HspB1_77-171_ variants were analyzed at a concentration of 10 μM in 200 mM ammonium acetate pH 6.9. Spectra of FLNC_d18-21_ and peptide titrations were recorded with settings: capillary 1.50kV; sample cone 30V; extractor cone 3V; backing pressure 3.8 mbar; trap gas (argon) 4 mL min^−1^; trap and transfer cell voltages 5V. FLNC concentration was held constant at 5 μM and peptide concentration (HspB1_80-88_ or ^P^HspB1_80-88_) was varied from 5 to 160 μM. Peptides and protein were mixed in 200 mM ammonium acetate pH 6.9 immediately prior to data collection. Nanoelectrospray ionization was performed in the presence of acetonitrile vapor in order to promote charge reduction and maintenance of noncovalent contacts and native structure (117). A minimum of three technical repeats were collected at each ratio. Dissociation constants were obtained by fitting the data using UniDec (118). To control for nonspecific adduct formation during desolvation, spectra were collected using an excess of PPR peptide with scrambled sequence. To control for the possibility of promiscuity in recognition of Ig-domain folds by HspB1 flexible termini, spectra were collected using an excess of peptide mimicking HspB1 C-terminal residues 177-186 (Fig. S5).

Gas-phase unfolding experiments were performed using the same conditions as described for peptide titrations, with an ion mobility cell gas pressure of 0.53 mbar (N_2_) and flow rate 22 mL min^−1^, at wave velocity 300 m s^−1^ and wave height 7.0V. Collision voltage into the trap cell was varied from 5 to l80 V, in 5-V increments below 100 V, and 10-V increments above 100 V. All data shown herein are from unfolding experiments performed with FLNC in the presence of 16-fold molar excess of peptide, excepting controls in Fig. S7. Plots are representative of experiments performed with three different protein batch preparations and three peptide to protein ratios. To quantify peptide dissociation with activation, the 13+ singly bound peak was selected in the quadrupole with LM and HM resolution 7.2 and 6.9, respectively, and trap voltage ramped as described for the unfolding experiments. Data were smoothed 30x2 in MassLynx and peak intensities recorded, summed as bound or unbound, and converted to percentages. A subtraction was applied to correct for nonspecific adduction at potentials of 30V and below based on activation trials with nonspecific controls.

### Calculation of experimental and predicted collision cross-sections

CCSs of models from the PDB (trimmed in PyMol as described in Supplementary Materials) were calculated by the projection approximation method using IMPACT (119) accounting for N_2_ as the collision gas. To calculate CCSs of FLNC_d18-21_, native mass spectra of β-lactoglobulin and avidin were collected under identical drift cell conditions (as described for titration experiments, with the mobility wave height lowered to 5.5 V) and used to construct a calibration in PULSAR (120) which had R^2^>0.96, using a power law linear fit.

### SDS-PAGE aggregation assay

HspB1 (WT or 3P) was mixed with FLNC_d18-21_ at a 2 to 1 ratio (80 and 40 μM, respectively) in 200 mM ammonium acetate pH 6.9 with 500 μM DTT and incubated at 25°C for 90 minutes. Aliquots were removed periodically and pelleted at 13,000 g at 10°C for 10 minutes. Soluble fractions were diluted three-fold with 200 mM ammonium acetate for analysis by SDS-PAGE, and pellets were resuspended in the same volume of 200 mM ammonium acetate and solubilized upon addition of the SDS-PAGE loading buffer. Gel bands were quantified by densitometry, using the raw intensities of FLNC and HspB1 bands to extract relative soluble and insoluble proportions at each time point.

### MD simulations

Two simulations of the HspB1 dimer were prepared using a previous ACD crystal structure (PDB: 4MJH). The system was immersed into a TIP3P water box, and neutralized with 0.15 M Na+ and Cl− ions. Simulations used the Amber99SB force field (121) on NAMD2.9 (122) molecular dynamics engine, with SHAKE algorithm constraining heavy atoms distances, and PME treating the electrostatic interactions in periodic boundary conditions. The systems were first minimized with 2000 conjugate gradient steps, and subsequently simulated using a 2 fs timestep. Systems were equilibrated in the nPT ensemble for 0.5 ns, with a 10 kcal mol^−1^ constrain on protein alpha carbons. 300 K and 1 atm were imposed by Langevin dynamics, using a damping constant of 1 ps^−1^, a piston period of 200 fs, and a piston decay of 50 fs. Constraints were subsequently removed, and systems were simulated in the nVT ensemble at 300K for 1 ns. Finally, production runs were conducted in the nPT ensemble (300K and 1 atm) for 1 μs. We extracted the coordinates of each individual monomer every 0.1 ns, yielding 1000 frames per monomer. We calculated the order parameter (*S*^*2*^ of N-H bond vectors) of the datasets independently with a Python code developed in house.

## Acknowledgments

We thank Stephen Weeks and Sergei Strelkov (KU Leuven) for providing the SUMO-HspB1 plasmid. M.P.C. was supported by the Clarendon Fund (Oxford University Press), T.R.A. by Pembroke College and the NIH Oxford-Cambridge Scholars Program, and H.Y.G. by the Doctoral Training Centre in Systems Biology. We thank the Biotechnology and Biological Sciences Research Council (J.L.P.B. for BB/K004247/1 and BB/j018082/1), and Diamond Light Source for access to beamline I04-1 (MX12346). K.D-C. research is supported by the Austrian Science Fund (FWF) Projects I525 and I1593, P22276 and W1221, Federal Ministry of Economy, Family and Youth through the initiative “Laura Bassi Centres of Expertise”, the Centre of Optimized Structural Studies, No253275, and by the Welcome Trust Collaborative Award (201543/Z/16/Z). K.G. is supported by the British Heart Foundation (BHF; FS/12/40/29712), the BHF Centre of Research Excellence Oxford (RE/13/1/30181), and the Wellcome Trust (201543/B/16/Z).

## Author Contributions

Conceptualization - M.P.C., T.R.A., C.dV., D.O.F., P.vdV., K. D-C., K.G., J.L.P.B. Investigation - M.P.C., T.R.A., C.dV., D.N., H.Y.G., T.M.A., M.T.D., K.G. Writing (original draft) - M.P.C., J.L.P.B. Writing (review and editing) - all authors. The authors declare no conflict of interest.

